# Spontaneous locus coeruleus bursts coincide with transient global brain state changes similar to those elicited by surprise

**DOI:** 10.64898/2026.05.19.726177

**Authors:** Richard Somervail, Mingyu Yang, Giandomenico Iannetti, Oxana Eschenko

**Affiliations:** Translational and Computational Neuroscience Unit, Manchester Metropolitan University, Manchester, M15 6GX, State, United Kingdom; Neuroscience and Behaviour Laboratory, Italian Institute of Technology, Rome, 00161, State, Italy; Department of Neuroscience, Physiology and Pharmacology, University College London, London, WC1E 6BT, State, United Kingdom; Department of Physiology of Cognitive Processes, Max-Planck Institute for Biological Cybernetics, Tübingen, 72076, State, Germany; Department of of Physiology and Pharmacology,, Rome, 00185, State, Italy; Department of Body-Brain Cybernetics, Max-Planck Institute for Biological Cybernetics, Tübingen, 72076, State, Germany

**Author notes:** Corresponding authors, shared senior authorship.

**Keywords:** noradrenaline, neuromodulation, phasic response, extralemniscal system, sensory processing

## Abstract

Sudden and isolated sensory stimuli (SISS) engage the extralemniscal system and elicit widespread electrocortical responses in the brain. These responses, consisting of both time-domain transients and spectral changes, reflect a switch of the global brain state that likely prepares the organism for subsequent urgent behaviours. Crucially, SISS also elicit a short-latency phasic response in a key component of the extralemniscal system in the brainstem, the noradrenergic Locus Coeruleus (LC) nucleus. Such stimulus-evoked LC firing is associated with the electrocortical markers of extralemniscal activation. LC neurons also display burst-like firing spontaneously, i.e., without imposed sensory stimuli, for example, during quiet wakefulness, sleep, or anaesthesia. However, this phenomenon remains underexplored. We therefore measured, in freely behaving rats, the prefrontal electrocorticogram (ECoG) responses following spontaneous LC bursts. In addition, we compared these ECoG responses to those triggered by electrical LC stimulation or auditory SISS. We found that ECoG responses were proportional to the magnitude of the spontaneous LC bursts or microstimulation, and remarkably similar to those elicited by SISS. Finally, suppression of noradrenergic transmission with systemic clonidine administration attenuated the auditory-evoked ECoG response. These results suggest that LC plays a role in generating the transient brain state changes elicited by SISS.

## Introduction

Survival in dynamic environments depends on rapid, adaptive behavioural responses. An important computational challenge that the central nervous system (CNS) must solve is to determine which environmental changes are relevant to ongoing behaviour. Accordingly, the CNS is particularly attuned to detect sudden and unexpected events such as the appearance of a predator, an insect sting, or an opportunity to catch prey. Such events typically trigger sympathetic “fight-or-flight” responses associated with swift behavioural reorganisation.

These sudden and isolated sensory stimuli (SISS) elicit large and widespread neural responses, detected in population-level electrocortical recordings (EEG and ECoG) as “vertex potentials (VPs)” (Davis, 1939; Bancaud et al., 1953; Davis and Zerlin, 1966; Mouraux and Iannetti, 2009; Valentini et al., 2011; Torta et al., 2012; Ronga et al., 2013; Moayedi et al., 2016; Novembre et al., 2018; Somervail et al., 2021, 2022); for a review see (Somervail et al., 2025). The VP is accompanied by characteristic spectral changes, namely by increased high-frequency and decreased low-frequency cortical activity, a common index of an increase of arousal (Mouraux et al., 2003; Tiemann et al., 2010; Zhang et al., 2012; Peng et al., 2018; Hu and Iannetti, 2019). These electrophysiological markers reflect a change of the global brain state, likely allowing the rapid and flexible reallocation of cognitive resources towards the preparation of appropriate behaviours (Somervail et al., 2025).

We have recently highlighted the crucial role of the non-specific “extralemniscal” sensory system in mediating this change of brain state (Somervail et al., 2025). The extralemniscal system includes diffuse thalamocortical projections and the brainstem reticular formation (Abrahamian et al., 1963; Jones, 2003; Bellesi et al., 2014; Whyte et al., 2024), which modulate various dimensions of the central arousal state (De Gee et al., 2020; Munn et al., 2021). The brainstem noradrenergic nucleus Locus Coeruleus (LC) is a key component of the reticular ascending arousal system (Berridge, 2008; Sara and Bouret, 2012; Ross and Van Bockstaele, 2021). SISS elicit phasic discharge of LC neurons (Foote et al., 1980; Aston-Jones and Bloom, 1981b; Hurley et al., 2004; Edeline et al., 2011; Martins and Froemke, 2015; Glennon et al., 2019; McBurney-Lin et al., 2020). In our previous studies, we have shown that direct microstimulation of LC mimics sensory-evoked LC bursting, resulting in immediate increases of cortical arousal lasting several seconds and evokes large-amplitude transients and spectral changes in the EEG (Marzo et al., 2014; Neves et al., 2018; Yang et al., 2021). Importantly, during sleep (Eschenko et al., 2012) or anaesthesia (Totah et al., 2021), LC neurons also display burst-like firing spontaneously, i.e., in the absence of imposed sensory stimuli. Interestingly, spontaneous LC bursts, occurring under anaesthesia, were associated with enhancements of high-frequency prefrontal activity - a cortical response reminiscent of that observed in response to noxious somatosensory stimuli (Neves et al., 2018). However, this phenomenon has not been studied in behaving animals, where the focus has been mainly on sensory-driven LC phasic responses. As such, cortical responses to spontaneous phasic LC activity remain unexplored.

Therefore, here we investigated spontaneous LC bursting in wakefulness and its relationship to changes in the global brain state as revealed by electrocortical activity. Specifically, we hypothesized that LC burst firing might contribute to the generation of the same ECoG response as the one elicited by SISS. We therefore analysed prefrontal ECoG signals in behaving rats during quiet wakefulness. We measured the temporal and spectral profiles of the ECoG responses following spontaneous LC bursts occurring in the absence of experimentally controlled sensory input. We also compared the ECoG responses associated with spontaneous LC bursts to those triggered by direct electrical LC stimulation or external auditory SISS. Finally, to provide causal evidence for the role of the LC, we pharmacologically suppressed noradrenergic transmission by systemic administration of clonidine while delivering auditory SISS.

## Materials and Methods

### Animals

We reanalysed the data that were collected for our previous studies (Eschenko et al., 2012; Novitskaya et al., 2016; Yang et al., 2021; Yang and Eschenko, 2025). The datasets from 21 adult male Sprague-Dawley rats (RRID:RGD 70508), weighing 300 - 350 g at the beginning of the experiment, were included for analyses (Supplementary Table 1). Briefly, rats (Charles River Laboratory, Germany) were kept on a 12-hour light-dark cycle (8:00 am lights on) and single-housed after surgery. Recordings were performed during the dark cycle. Similar surgery and recording procedures were used as reported in detail elsewhere (Eschenko et al., 2012; Yang et al., 2021; Yang and Eschenko, 2025). All experimental procedures were performed in accordance with the German Animal Welfare Act (TierSchG) and Animal Welfare Laboratory Animal Ordinance (TierSchVersV). This is in full compliance with the guidelines of the EU Directive on the protection of animals used for scientific purposes (2010/63/EU). The study was reviewed by the ethics commission (§15 TierSchG) and approved by the state authority (Regierungspräsidium, Tübingen, Baden-Württemberg, Germany).

### Surgery and electrode placement

Procedures for stereotaxic surgery under isoflurane anesthesia have been described in detail elsewhere (Yang et al., 2021; Yang and Eschenko, 2025). Briefly, anesthesia was initiated with 4% and maintained with 1.5-2.0% isoflurane. Body temperature, heart rate, and blood oxygenation were monitored throughout the entire anesthesia period. The depth of anesthesia was controlled by a lack of pain and sensory response (hind paw pinch). A fully anesthetized rat was fixed in a stereotaxic frame with the head angle at zero degrees. The skull was exposed, and a local anesthetic (Lidocard 2%, B. Braun, Melsungen, Germany) was applied to the skin edges. Craniotomies were performed on the right hemisphere above the target regions. Additional burr holes were made for EEG, grown, and anchor screws (stainless steel, 0.86 - 1.19 mm diameter, Fine Science Tools, Heidelberg, Germany). Dura mater was removed when necessary. For LC recording, single platinum-iridium electrodes (FHC, Bowdoin, ME, n = 1), microwires brush electrode (MicroProbes, MD, n = 2), and a silicon probe (Cambridge Neurotech, Cambridge, UK, n = 2) mounted on a microdrive (Cambridge Neurotech, Cambridge, UK) were implanted above LC (*∼* 4.2 mm /*∼* 1.2 mm from Lamda; DV: *∼* 5.5 - 6.2 mm) at a 15-degree angle. The accuracy of LC targeting was verified by online monitoring of neural activity. The LC neurons were identified by broad spike widths (*∼* 0.6 ms), regular low firing rate (1 – 2 spikes/s), and a biphasic (excitation followed by inhibition) response to paw pinch. For EEG recording, a screw was placed above the frontal cortex. The ground screw was placed above the cerebellum.

Screws were fixed in the skull and additionally secured with tissue adhesive. The entire implant was secured on the skull with dental cement (RelyX™ Unicem 2 Automix, 3M, MN). A copper mesh was mounted around the implant for shielding and protection of exposed connection wires. During the post-surgery recovery period, analgesic (2.5 mg/kg, s.c.; Rimadyl, Zoetis) and antibiotic (5.0 mg/kg, s.c.; Baytril®) were given for 3 and 5 days, respectively. Electrode placements were histologically verified.

### Experimental Design

Sudden and intense auditory stimuli were presented to spontaneously behaving rats (quiet awake/sleep state) at randomized intervals of 10-20 s. 40-ms tones of 65-105 dB (with 5dB steps) were presented in random order on the background of white noise (50/55 dB; (Yang et al., 2021). Electrical LC microstimulation consisted of trains of biphasic (cathodal leading) square pulses (0.4 ms, 0.05 mA) presented at 20, 50 and 100 Hz for 100 – 200 ms (Novitskaya et al., 2016).

### Drugs

Clonidine was obtained from Sigma-Aldrich (Deisenhofen, Germany). Before each recording session, rats received the intraperitoneal (i.p.) injection of clonidine (*α*2-adrenergic receptor agonist, 0.05 mg/kg).

### Electrophysiological recording and data analysis

Rats were first habituated to the sleeping box (40 cm height, 30 cm diameter) and cable plugging procedure. The electrodes were either connected to the Neuralynx Digital Lynx or FreeLynx wireless acquisition system via two 32-channel head stages (Neuralynx, Bozeman, MT). The electrode placement in the LC was optimized by lowering the electrodes with a maximal 0.05 mm step and monitoring spiking activity on the high-passed (300 Hz – 8 kHz) extracellular signal. Once the electrode position was optimized, the broadband (0.1 Hz - 8 kHz) extracellular signals were acquired and digitized at 32 kHz and referenced to the ground screw. The animals’ movement was monitored by an EMG electrode attached to the neck muscle. All recordings were performed between 10 a.m. and 7 p.m.

ECoG signals were imported into EEGLAB (RRID:SCR 007292) (Delorme and Makeig, 2004), band-pass filtered between 1 and 100 Hz, and resampled to 500 Hz. Continuous recordings were epoched *±*2 s around the trigger event, depending on the experimental condition.

### ECoG Peak Analysis

For the saline and clonidine data, we extracted single-subject peak amplitudes and latencies for each condition. Peaks were extracted from the low-pass filtered (Butterworth, cutoff = 50 Hz, order = 4) single-subject averaged waveforms for each condition and quantified as the most negative data point in the window from 0 to 100 ms. These amplitude and latency values were then compared between the saline and clonidine conditions with paired t-tests.

### ECoG time-frequency analysis

The epoched ECoG time-domain signals were transformed to the time-frequency domain using a wavelet analysis (EEGLAB, *pop newtimef*) with an 80-point linear frequency scale (5 - 100 Hz) and a sliding window (window interval = 20 ms). Amplitude values were computed as the absolute of the complex-valued convolution of the wavelets with the time-domain data. Inter-trial phase-coherence (ITPC) values were also computed. Amplitude and ITPC values per frequency band were then baseline corrected by trial-wise subtraction of the mean value from -1 to -0.5 s.

When comparing the high-frequency ECoG amplitude associated with high- and low-frequency LC bursts (Figure 2) or auditory stimuli with or without the presence of clonidine (Figure 6), we performed an additional ROI analysis to summarise the relatively sparse and low-amplitude high-frequency activity, by computing the mean values in an ROI covering the high frequency range (30 - 100 Hz) in the time-window: 0.1 - 1 s (Figures 2 and 6, dashed box on time-frequency plots).

### Classification of behavioural states

We classified the rat’s spontaneous behaviour into awake and NREM sleep using frontal EEG or cortical LFPs by applying a standard sleep scoring algorithm described in detail elsewhere (Novitskaya et al., 2016; Yang and Eschenko, 2025). Briefly, animal movement speed was extracted from the video recording synchronized with a neural signal. The theta/delta ratio (*θ*/*δ*) was calculated from the artifact-free EEG in 2.5-sec epochs. The epochs of the awake state were identified by the presence of active locomotion and above *θ*/*δ* threshold; the epochs of NREM sleep were identified by the absence of motor activity and below *θ*/*δ* threshold. The minimal duration of the same behavioural state was set to 20 sec; data segments with less steady behavioural states were excluded from further analysis.

### LC burst detection

Spike-density functions (SDF) were computed for all LC recordings by convolution of the detected spikes with a Gaussian kernel (SD = 60 ms), resulting in an SDF time-series sampled at 1000 Hz. These SDF values were then imported into EEGLAB (Delorme and Makeig, 2004) for subsequent analysis. LC bursts were detected by the following method: First, the spike-density function (SDF) of the LC channel containing multi-unit spike trains was computed by convolution of the spike vector with a Gaussian kernel (SD: 60 ms). Second, the SDF was z-scored (mean subtracted and divided by its SD). Third, this z-scored SDF was used to identify burst peaks as local maxima with a magnitude larger than 3 SD, separated from each other by at least 1 s. Finally, the onset of each burst was identified by finding the first spike that arrived later than its predecessor by more than 30 ms. If no preceding spike was found within a 0.5 s window, then the burst onset was defined as the time point 0.5 s before the peak in the SDF.

Once LC bursts were detected, we found the corresponding bursts on the raw SDF (before z-scoring), epoched the signals between -2 and +2 relative to the estimated burst onsets, and then baseline corrected each epoch by subtracting the mean SDF across the -2 to -1 s pre-burst window concatenated across all epochs. This approach allowed us to quantify the LC burst magnitude as a proxy of phasic NA release, while controlling for tonic LC firing.

Only auditory stimuli with intensities above 90 dB SPL were retained for further analysis. In all conditions, epochs were rejected if the ECoG signal was (1) flat throughout the epoch or (2) higher than 1 mV. Supplementary Table 2 shows, for each experimental condition, the mean and SD of the total number and percentage of rejected trials across subjects.

### LC burst median split analysis

To compare the ECoG responses in trials with larger or smaller LC bursts (Figure 2), we performed a median split analysis, in which we computed the trial-wise LC burst magnitudes as the most positive local maxima in the time-window from -0.05 to 0.3 s relative to burst onset (using MATLAB’s *findpeaks* function). Trials whose peak value was larger than the median value were considered “high-frequency” bursts, while those equal to or less than the median were considered “low-frequency” bursts.

### LC burst-matching procedure

In the single-animal analysis shown in Figure 5, we selected a subset of LC bursts that arose spontaneously and those associated with auditory stimuli in which their peak frequencies were approximately equal. This procedure involved (1) measuring the peak frequency of each baseline-corrected LC burst in both conditions, (2) automatically separating these peak frequencies into 5 evenly-spaced bins, and (3) selecting the subset of bursts in each condition whose peak frequency fell within the second bin (to achieve a balance between sample size and burst frequency).

### Statistical Analysis

The consistency of group-level mean responses to all conditions was statistically assessed by one-sample two-tailed t-tests against zero (MATLAB). All plots show 95% confidence intervals resulting from these t-tests. The difference between the group-level mean responses of different conditions was assessed with paired or unpaired t-tests depending on the conditions (see Results). T-tests were all run separately for each time point and separately for the high-frequency ROI analysis. The resulting P values from these tests were rescaled to surprisal (S) values by computing their negative base-2 logarithm (S = -log2(P); (Greenland, 2017). Surprisal values provide an intuitive binomial scale for the degrees of evidence expressed by P values. They can be interpreted as the number of coin flips landing on ‘heads’ in a row (with the typical P = 0.05 threshold corresponds to a surprisal value of *∼*4.3 bits).

## Results

We simultaneously recorded extracellular spiking of putative LC-NA neurons and prefrontal ECoG in freely behaving adult male rats (n = 5) in the absence of any experimental sensory stimulation (see Methods for details). Recordings lasted between 31.9 and 175.1 minutes (mean *±* SD: 92.8 *±* 26.2 mins) without interruptions. During recordings, the rats’ spontaneous behavior typically alternated between wakefulness (18.2 - 100% recording time; 55.3 *±* 22%) and sleep (0 - 81.8% recording time; 44.7 *±* 22%). We focused our analysis on the wakefulness episodes.

### Spontaneous LC bursting is reflected in electrocortical activity

In quiet wakefulness, LC neurons displayed irregular synchronized firing (“LC bursts”) in the absence of sensory stimulation. To identify LC bursts, we first generated a spike density function (SDF) by convolving the multi-unit spike train with a Gaussian kernel (SD = 60 ms). Bursts were then detected as local maxima with the SDF exceeding 3 SD and separated from other such maxima by at least 1 s. For each of these events, the burst onset was defined as the first spike that followed its predecessor by ¿ 30 ms. Spontaneous LC bursting varied from 2.2 to 18.6 bursts per minute (7.4 *±* 5.2). Figure 1A shows an example raster plot from the detected LC bursts in a single session. Figure 1B shows the group-level average SDF of the detected LC bursts (N = 5). Figure 1C and 1D show the changes in the ECoG associated with LC bursts in the time and time-frequency domains, respectively. Figure 1E shows phase-locking values of the time-frequency ECoG response (inter-trial phase coherence; ITPC). Spontaneous LC bursts were associated with consistent large-amplitude negative deflections of the time domain ECoG (Figure 1C). This time-domain deflection corresponded, in the time-frequency domain, to the bottom part (i.e., *∼* 5 - 7 Hz) of the low-frequency, early-latency amplitude increase, which was relatively phase-locked to burst onset (Figure 1D and 1E). This early latency response coincided with a non-phase-locked high-frequency (20 - 100 Hz) amplitude increase starting at *∼* 0.1 s, and was followed by a sustained low-frequency (5 - 20 Hz) amplitude decrease starting at *∼* 0.2 s, both of which lasted up to 1 second after burst onset (Figure 1D and 1E).

**Figure 1.**
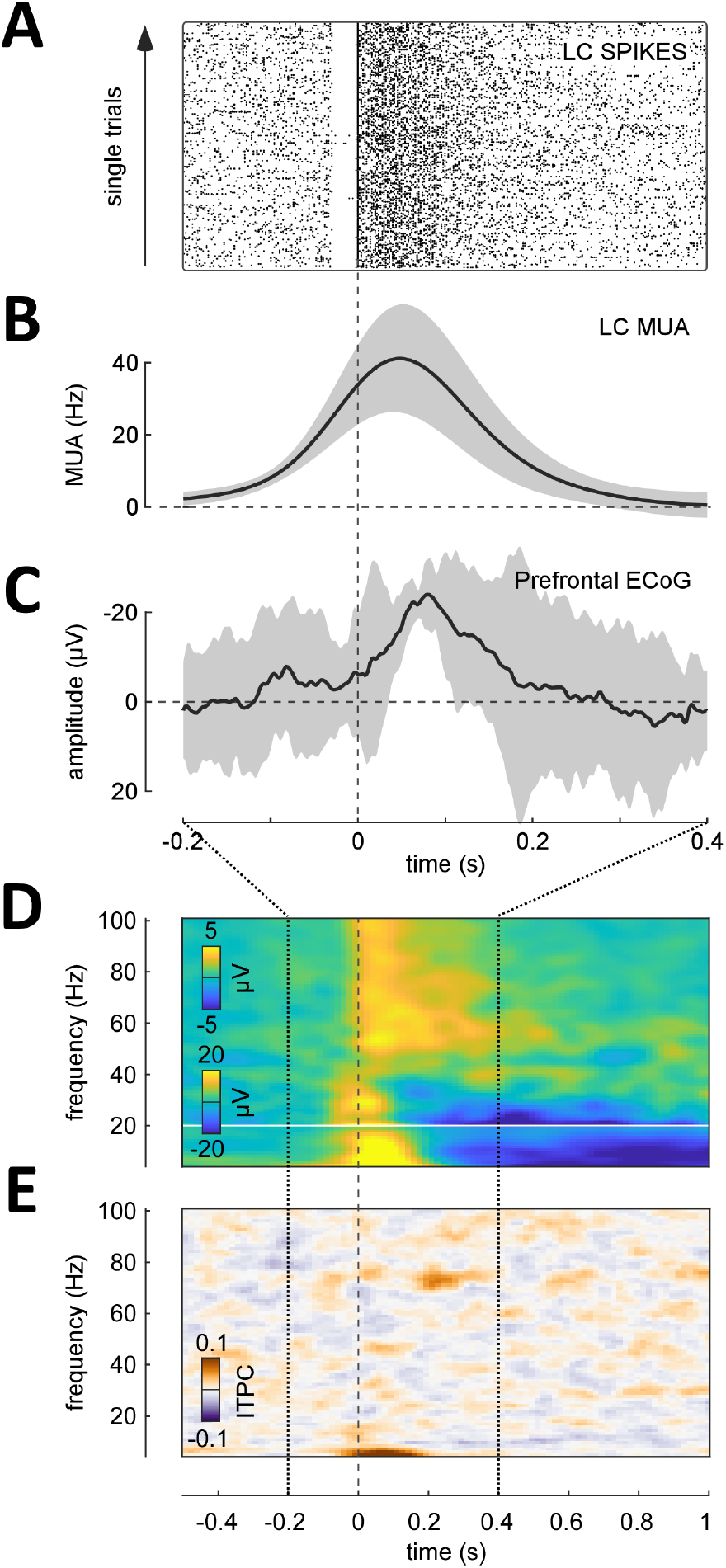
Cortical response is time-locked to spontaneous bursting of locus coeruleus (LC) neurons during wakefulness in the absence of external stimuli. **(A)** Raster plot of spontaneous LC bursting from a representative animal. Bursts were detected using an automatic approach (see Methods). Rows represent trials sorted by their order of occurrence, aligned according to burst onset (time = 0 s). **(B)** The group-averaged spike-density function of the LC-NA spikes (Multi-Unit Activity, MUA) is plotted around the onset of the detected LC bursts. Shaded area indicates 95% confidence intervals computed with a t-test against zero. **(C**,**D)** Spontaneous phasic activation of the LC was associated with large-amplitude ECoG responses both in the time-domain **(C)** and in the time-frequency domain **(D)**. Note the two different colour scales to highlight amplitude changes of low (5 - 20 Hz) and high (20 - 100 Hz) frequencies. After the LC burst onset, there was a broadband transient amplitude increase (20 - 100 Hz), followed by a longer-lasting amplitude decrease at low frequencies and an amplitude increase at high frequencies. These spectral changes are known markers of cortical arousal (Yang et al., 2021). **(E)** Group-level average change of inter-trial phase-coherence (ITPC) associated with spontaneous LC burst. ITPC shows that most spectral changes shown in **D** were not phase-locked to the burst onset, except for the bottom part of the transient low-frequency amplitude increase.

### Larger spontaneous LC bursts are associated with larger cortical responses

After having ascertained that spontaneous LC bursting is associated with pronounced changes in the ECoG, we explored whether these cortical responses depended on the LC burst size. To this end, we extracted the peak frequency of each LC burst from the SDF and median split trials by intra-burst frequency (Figure 2A, *insert*). We then compared the ECoG changes associated with low- and high-frequency LC bursts. In the time domain, higher-frequency LC bursts corresponded to larger negative ECoG deflections (Figure 2A). In the time-frequency domain, both the amplitude and the duration of the ECoG spectral frequency modulation were proportional to the LC burst size (Figure 2B). Specifically, longer-lasting (0.2 - 1 s) power amplitude decreases at low-frequencies (5 - 20 Hz) were observed around bigger LC bursts. At the high-frequency range (30 - 100 Hz), there was sparse evidence of a higher-amplitude and longer-lasting (0.1 - 1 s) ECoG change associated with bigger LC bursts. Since this sparse effect was not well assessed by the point-by-point analysis used to obtain these results, we performed a paired t-test comparing the high- and low-frequency LC burst conditions using the mean values within an ROI covering this long-lasting increase of high-frequency amplitude (30 - 100 Hz, 0.1 - 1 s; Figure 2B, top panel). This analysis revealed evidence of a stronger and longer-lasting high-frequency amplitude increase around bigger LC bursts (mean difference = 1.31 *µ*V, d = 1.44, t_(4)_ = 3.22, P = 0.032, S = 4.96 bits).

**Figure 2.**
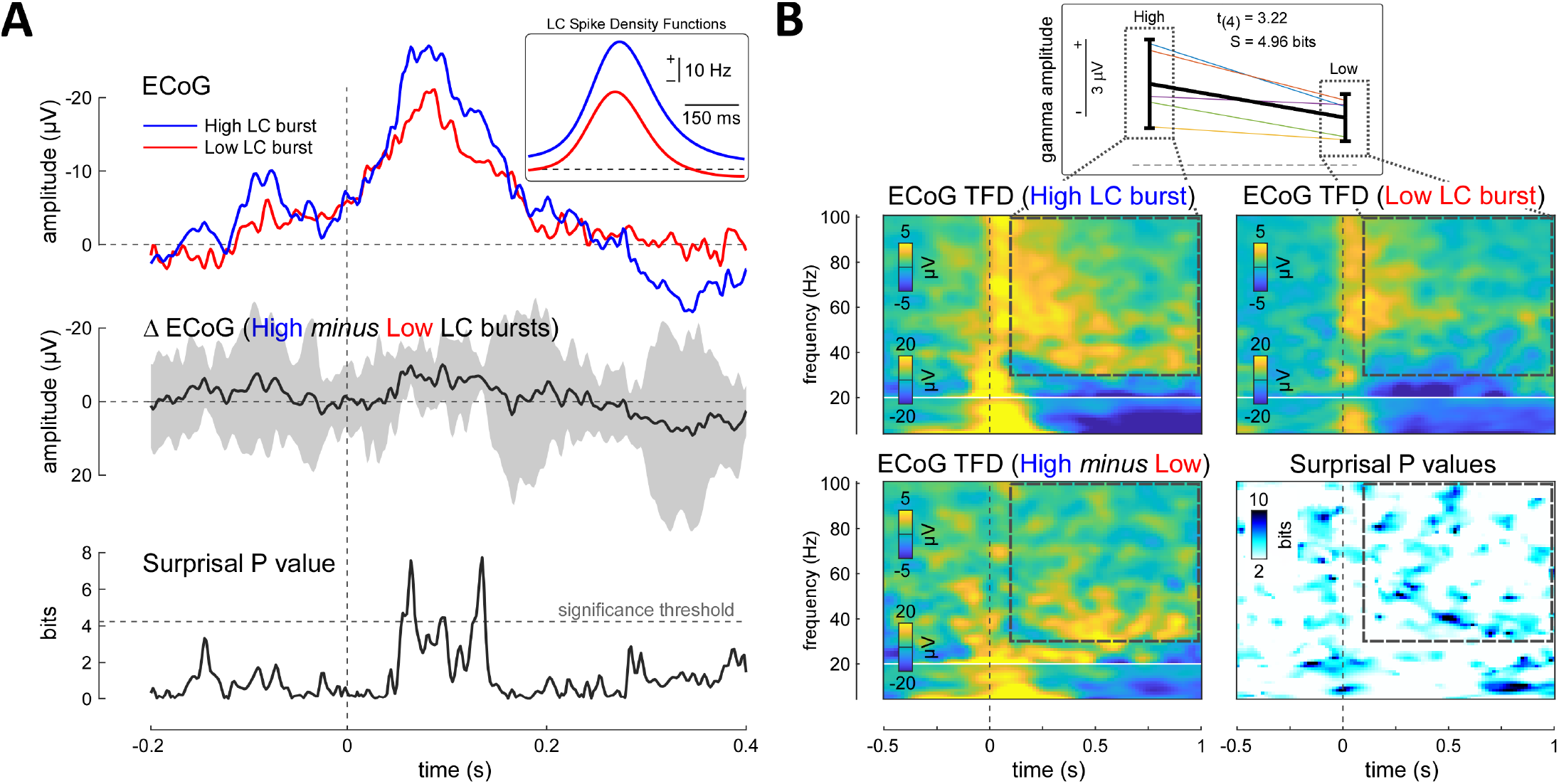
Higher-frequency LC bursts were associated with larger ECoG responses. All detected LC bursts were split within each animal according to whether their peak spike-density function (SDF) value was lower or greater than the median. **(A, top)** Grand average time domain ECoG responses associated with low (red) and high (blue) frequency LC bursts. **(A, middle)** Mean paired difference of the ECoG signals to low and high frequency LC bursts. The shaded area shows 95% confidence intervals from the point-by-point paired t-tests between the low and high burst conditions. **(A, bottom)** Surprisal values from the t-tests. Higher-frequency LC bursts resulted in a small but consistent increase of the time-domain ECoG response. **(B)** Grand average time-frequency domain ECoG signals (TFDs) associated with high (*top left*) and low (*top right*) frequency LC bursts, mean difference between these two conditions (*bottom left*), and the corresponding surprisal P values (*bottom right*). Note the two different colour scales to highlight amplitude changes of low (5 - 20 Hz) and high (20 - 100 Hz) frequencies. Both the initial transient increase and the subsequent long-lasting decrease (from 0.3 to 1 s) of low-frequency (5 - 20 Hz) amplitude were larger when elicited by higher-frequency spontaneous LC bursts. The top panel shows single-subject average amplitude values from an ROI including the high frequency response (30 - 100 Hz; 0.1 - 1 s). This analysis revealed clear evidence of a larger high-frequency amplitude increase associated with the high frequency compared to the low frequency LC bursts (paired t-test). Altogether, these results provide correlational evidence that these ECoG responses are consequent to the LC activation.

### Phasic LC microstimulation induces large-amplitude ECoG responses

We have previously demonstrated that phasic LC microstimulation mimics the activation of LC neurons induced by nociceptive stimuli in anaesthetized rats (Marzo et al., 2014). To compare the ECoG response to LC microstimulation and spontaneous LC bursting in freely behaving rats, we reused previously published data (Novitskaya et al., 2016). Specifically, in our previous study, we stimulated the LC with brief (100 ms) and intermittent (ISI: 10-20 s, uniform distribution) trains of electric pulses at 20, 50, and 100 Hz while simultaneously recording prefrontal activity (Novitskaya et al., 2016). The cortical responses elicited by LC microstimulation and associated with spontaneous LC burst showed striking similarities. Namely, they consisted of the typical large negative transient ECoG deflection in the time domain (Figure 3A), as well as the long-lasting increase of high-frequency and decrease of low-frequency amplitude (Figure 3B). Crucially, the amplitude of these cortical responses scaled with the frequency of LC microstimulation, with larger responses being evoked at higher stimulation frequencies (Figure 3).

**Figure 3.**
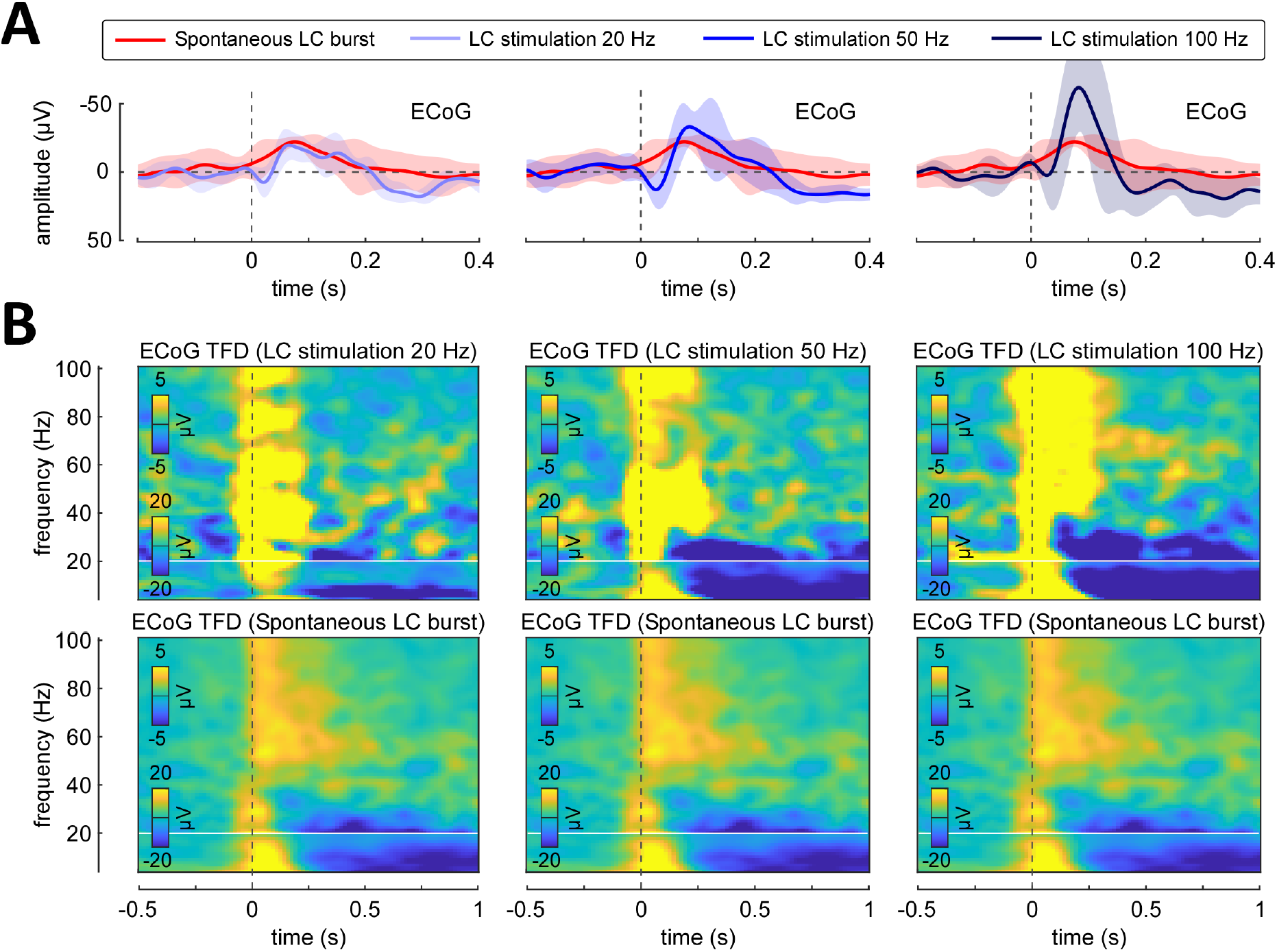
Phasic LC microstimulation evokes ECoG responses similar to those associated with spontaneous LC bursting. **(A)** Average peristimulus ECoG response (blue, low-pass filtered at 15 Hz) elicited by LC microstimulation at 20 Hz (left), 50 Hz (middle), and 100 Hz (right) in awake, free-behaving animals. For comparison, the ECoG responses associated with spontaneous LC bursts are shown in red. The time-domain response to LC stimulation was qualitatively similar to that associated with spontaneous LC bursts, but varied in amplitude depending on LC stimulation frequency. The shaded area indicates 95% confidence intervals computed with a t-test against zero. **(B)** The group-level time-frequency domain responses to LC microstimulation (top rows) and those associated with spontaneous LC bursts (bottom rows). Note the two different colour scales to highlight amplitude changes of low (5 - 20 Hz) and high (20 - 100 Hz) frequencies. The low-frequency transient increase of amplitude (from *∼* -0.05 to 0.2 s) and the longer-lasting decrease of low-frequency amplitude (from *∼* 0.2 s to 1 s) were similar in the two types of LC activation. Higher frequency LC stimulation produced overall stronger spectral modulations. Thus, direct LC stimulation elicits an ECoG response highly similar to those associated with spontaneous LC bursting, providing direct causal evidence for the LC’s involvement in generating this ECoG response. Note that the broadband (5 - 100 Hz) transient response in the LC stimulation condition is contaminated by electrical stimulation artifacts and is therefore not clearly visible.

### Cortical responses associated with spontaneous LC bursts resemble auditory-evoked but are less robust

At a cursory glance, the ECoG changes associated with spontaneous LC bursts were reminiscent of the human EEG responses to SISS, which are mediated by a transient activation of the extralemniscal system (Mouraux et al., 2003; Mouraux and Iannetti, 2009; Hu and Iannetti, 2019; Somervail et al., 2021). To formally explore this similarity within-species, we analysed ECoG recorded while rats were exposed to auditory SISS. We used previously published data (Yang et al., 2021) and selected trials with the sound intensity 90 -110 dB (Supplementary Table 1), which reliably produced an overt behavioral startle response (Yang et al., 2021). Figure 4 shows auditory SISS-evoked ECoG responses both in the time and time-frequency domains. The auditory SISS elicited a large-amplitude time-domain ECoG transient, reminiscent of the bipolar VP in humans (Mouraux and Iannetti, 2009; Somervail et al., 2021), characterised by a large negative deflection peaking at *∼*54 ms, followed by a smaller positive deflection at *∼*132 ms. This negativity appeared to last slightly longer than the negativity associated with spontaneous LC bursts, but the presence of the subsequent positive deflection in the auditory-evoked response may have obscured the latter part of this potential. The time-frequency response to the auditory SISS comprised an early-latency phase-locked increase of broadband amplitude (5 - 100 Hz, -0.1 - 0.16 s; whose lower-frequency component reflects the time domain transient), followed by a longer-lasting decrease of low-frequency amplitude (5 - 20 Hz, *∼*0.2 - 1 s) and an increase of high-frequency amplitude (30 - 100 Hz, *∼*0.1 - 1 s). These ECoG signals were also reminiscent of those associated with spontaneous LC burst (Figure 1).

**Figure 4.**
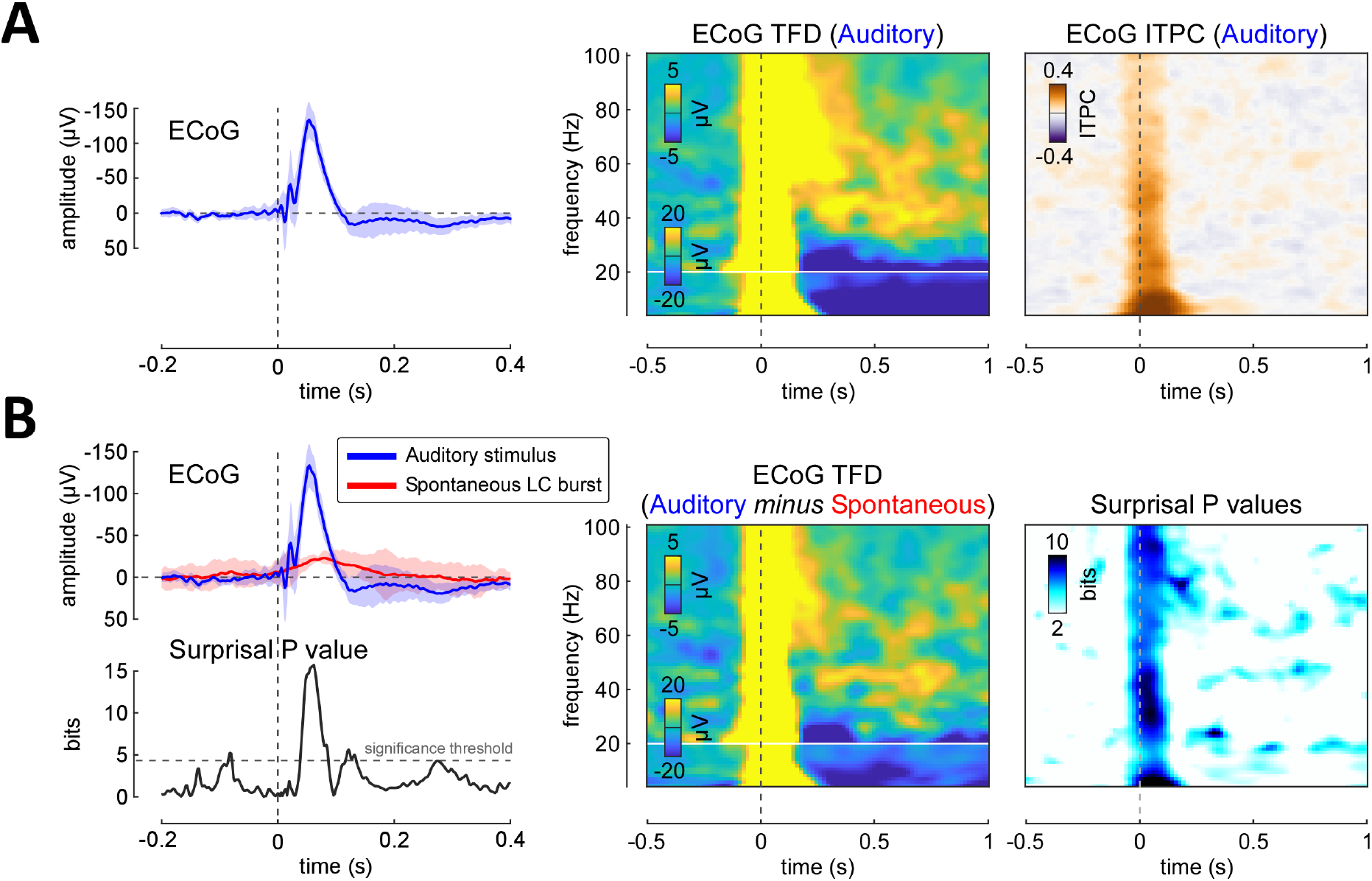
Comparison between ECoG responses to spontaneous LC bursting and auditory stimulation. **(A)** Auditory-evoked responses in the time domain (*left*), time-frequency domain (*middle*), and inter-trial phase-coherence (*right*). The shaded area in the left plot indicates 95% confidence intervals computed with a t-test against zero. **(B)** Unpaired t-test comparison of auditory ECoG response to that associated with spontaneous LC burst in the time-domain (*left*) and time-frequency domain (*middle*: difference plot, *right*: surprisal P values). Shaded areas indicate 95% confidence intervals computed with t-tests against zero. Note the two different colour scales to highlight amplitude changes of low (5 - 20 Hz) and high (20 - 100 Hz) frequencies. These results show that auditory stimuli elicit a time-domain ECoG response substantially larger than that associated with spontaneous LC burst, but that both responses consist of a main negative deflection. The auditory time-frequency ECoG response was qualitatively similar to the response to spontaneous LC bursts, but had a higher transient increase of phase-locked broadband amplitude (5 - 100 Hz; partly reflecting the time domain deflections), and a stronger high-frequency amplitude increase following the transient (*∼*0.1 s - 0.25 s, 30 - 100 Hz), indicating a larger increase of central arousal.

We quantitatively compared the time and time-frequency domain ECoG signals elicited by auditory SISS with those associated with spontaneous phasic LC activation using unpaired point-by-point t-tests (Figure 4B). This analysis showed strong evidence that the ECoG responses to auditory SISS were consistently larger in amplitude compared to those associated with spontaneous LC bursting both in the time domain (surprisal values above 15 bits in a time interval centered at *∼*50 ms post-stimulus) and in the early amplitude increases in the time-frequency domain (surprisal values above 10 bits at 5 - 100 Hz, *∼* -0.05 - 0.1 s). We also found sparse evidence for a stronger later increase of high-frequency (30 - 100 Hz, 0.1 - 1 s) and decrease of low-frequency (5 - 20 Hz, 0.1 - 1 s) amplitude in the auditory-evoked response. Overall, these findings indicate that spontaneous and evoked LC bursting elicit comparable cortical responses, with differences mainly in magnitude.

### Within-subject comparison of spontaneous and auditory-evoked LC bursts and ECoG signals

In the previous Results section, we described that auditory SISS elicit qualitatively similar, yet larger ECoG responses than those associated with spontaneous LC bursts. We explored this difference in magnitude directly using a single dataset where auditory SISS were delivered while recording LC spikes and ECoG simultaneously. Figure 5A shows the LC spiking elicited by auditory SISS and during spontaneous bursting, as well as the associated ECoG. The auditory-evoked LC response was slightly but consistently more robust than LC spiking during spontaneous bursting. In contrast to this relatively small difference in the LC firing pattern, the difference in ECoG responses to auditory SISS and associated with spontaneous LC bursts was much more pronounced, with earlier onset and larger ECoG changes elicited by auditory stimulation.

We reasoned that some of this difference could be consequent to the larger LC spiking activity in response to auditory stimulation. To account for this, we matched the frequency of LC spiking in the two conditions (i.e., spontaneous and stimulus-evoked bursting; see Methods). When comparing spontaneous and evoked LC bursts of similar magnitude, we observed that the SDF in spontaneous bursting had a slower rise slope, a later peak, and was longer-lasting than in evoked bursting. Although this matching procedure reduced the difference between the ECoG responses to evoked and spontaneous LC bursting, there was still a consistent difference (t-test, surprisal value peak = *∼* 14.6 bits, i.e., P value = *∼* 4 × 10^-5^), with earlier-onset and larger cortical responses to auditory stimulation. The latency of the ECoG response peak associated with spontaneous LC bursts was also longer, mirroring the delayed spiking of the spontaneous LC bursting. These findings indicate that ECoG response magnitude is not solely determined by LC burst size, but instead reflects the recruitment of distributed circuits, likely including thalamocortical pathways within the extralemniscal system (Somervail et al., 2025).

### Pharmacological suppression of NA release attenuates auditory ECoG responses

The main results described so far are (1) the phenomenological similarity between the ECoG responses elicited by auditory SISS and those associated with spontaneous LC bursts, and (2) the tight relationship between the ECoG response amplitude and the LC burst size. We therefore hypothesised that the cortical response evoked by auditory SISS was, at least partly, mediated by the cortical NA release consequent to the LC phasic firing. We explicitly tested this hypothesis by recording auditory-evoked ECoG responses in a subset of 6 animals with and without systemic administration of clonidine, an agonist of presynaptic inhibitory *α*2-adrenergic receptors.

Figure 6A shows the auditory-evoked time-domain ECoG responses in the clonidine and saline conditions. In the presence of clonidine, the time-domain ECoG responses were reduced in amplitude compared to the saline condition, and the late negative deflection occurring between *∼*0.12 to 0.22 s post-stimulus was totally abolished. The point-by-point analysis suggested the possibility that the main negative ECoG deflection occurring at *∼*0.06 s post-stimulus was delayed in the clonidine condition, although this statistical approach cannot properly distinguish amplitude and latency differences across conditions. We therefore performed an additional analysis by measuring the peak amplitude and latency of the largest negative deflection in a time window ranging from 0 to 100 ms in each single-subject across-trial average waveform. These amplitude and latency values were then compared between the clonidine and saline conditions with paired t-tests. These t-tests provided evidence of a reduced response amplitude in the clonidine condition (mean difference = -33.7 *µ*V, d = -1.18, t_(5)_ = -2.90, P = 0.0339, S = 4.88 bits) and no clear evidence of a longer response latency in the clonidine condition (mean difference = +13.7 ms, d = 0.88, t_(5)_ = 2.14, P = 0.0849, S = 3.56 bits).

Figure 6B shows the comparison of the auditory-evoked time-frequency ECoG responses in the same two conditions, using paired t-tests for each frequency and time point. This analysis revealed strong evidence that clonidine resulted in smaller increases of low-frequency amplitude (0 - 0.2 s, 5 - 12 Hz), smaller increases of high-frequency amplitude (around 0 s, 30 - 100 Hz), and a disruption of the long-lasting decrease of low-frequency amplitude (*∼* 5 - 12 Hz). We also observed sparse evidence of a reduction in the long-lasting amplitude increase at high-frequency (0.1 - 1 s, 30 - 100 Hz). Since this effect was not well assessed by the point-by-point analysis, we used a paired t-test to compare the mean values over an ROI covering the high-frequency range (30 - 100 Hz) in the time-window 0.1 - 1 s in the saline and clonidine conditions

(Figure 6B, *top panel*). This ROI analysis revealed weak evidence of a reduced long-lasting high-frequency amplitude increase in the clonidine condition (mean difference = 2.54 *µ*V, d = 1.08, t_(5)_ = 2.66, P = 0.0451, S = 4.47 bits).

## Discussion

In this work we report, for the first time, the relationship between spontaneous LC bursting and cortical activity in behaving rats during quiet wakefulness. By simultaneously recording prefrontal ECoG and LC extracellular spikes, we provide evidence that synchronous discharge of LC neurons in the absence of experimentally-controlled sensory stimuli is associated with a large amplitude population-level cortical response, similar to the cortical vertex potential (VP) elicited by sudden and isolated sensory events (SISS) (Somervail et al., 2025). We compared the cortical responses associated with spontaneous, sound-evoked, and direct (microstimulation) LC phasic activation. In all three types of LC phasic activation, the time-domain ECoG response consisted of a similar large negative deflection (peaking at *∼* 54 ms). The ECoG spectral changes consisted of a sustained decrease of low-frequency amplitude (5 - 20 Hz, *∼* 0.2 - 1 s) and a more transient increase of high-frequency amplitude (30 - 100 Hz, *∼* 0.1 - 1 s). Finally, pharmacological suppression of NA transmission by clonidine attenuated the auditory-evoked ECoG response, providing causal evidence for a role of LC phasic activation in the generation of the cortical responses to SISS.

In the following discussion, we interpret these findings in a broader context of neural responses to surprising events - both external (exteroceptive) and internal (interoceptive) - and their modulation by the LC-NA system.

### LC burst contribution to ECoG spectral changes

Three pieces of evidence suggest that phasic LC bursts (and resulting NA release) influence cortical dynamics indexed by characteristic spectral changes in the ECoG. First, these spectral changes scaled with the magnitude of LC bursts (Figure 2). Second, they were attenuated when NA transmission was decreased by clonidine (Figure 6). Third, a similar ECoG response pattern was elicited by both direct electrical LC stimulation (Figure 3) and sudden sensory stimulation (Figure 4), i.e., experimental manipulations that elicit robust LC bursting.

We previously reported that LC microstimulation elicits similar spectral changes in the prefrontal cortex in rats under anaesthesia (Marzo et al., 2014) or during NREM sleep (Novitskaya et al., 2016). Phasic optogenetic LC stimulation also results in similar spectral changes in the orbitofrontal cortex (Wang et al., 2024); moreover, the sustained increase of gamma activity correlated with increased neuron firing across various cortical regions (Wang et al., 2024). Likewise, SISS not only elicit LC phasic response, but also similar spectral changes in the rat (Zhang et al., 2012; Peng et al., 2018; Yang et al., 2021) and human EEG (Mouraux et al., 2003; Tiemann et al., 2010; Hu and Iannetti, 2019). Remarkably, local pharmacological inactivation of LC completely eliminated the spectral changes in the prefrontal cortex elicited by noxious somatosensory stimuli in anaesthetised rats (Neves et al., 2018).

It is commonly accepted that multiple, large-scale cortical-subcortical networks regulate the brain state (Dringenberg and Olmstead, 2003), with LC-mediated widespread cortical spectral changes serving as a prototypical example of network-level neuromodulation. These LC-mediated changes of cortical activity could be explained by the direct and diffuse LC projections to the cortex (Constantinople and Bruno, 2011), non-specific thalamus Whyte et al. (2024), or indirectly via NA-mediated modulation of cholinergic transmission (España and Berridge, 2006). Indeed, high-frequency electrical stimulation of the basal forebrain (Goard and Dan, 2009) or non-specific thalamus (Hunter and Jasper, 1949; Jasper, 1949) also elicit spectral changes resembling the effects of LC phasic activation and enhanced arousal (Redinbaugh et al., 2020; Bastos et al., 2021; Tasserie et al., 2022; Zhang et al., 2024).

The pattern of the EEG spectral changes that we describe could therefore be a useful non-invasive biomarker of phasic LC activation in humans. One limitation is that other ascending neuromodulatory neurons also fire in bursts (Nuñez, 1996; Lin et al., 2006) and elicit similar spectral changes (Berlucchi, 1997; Dringenberg and Olmstead, 2003; Lin et al., 2006; Goard and Dan, 2009), so the resulting spectral changes can be considered a necessary but not sufficient condition to infer phasic LC activation. Moreover, noradrenergic innervation of cholinergic neurons in the basal forebrain suggest that the LC activity may contribute to the effects of the basal forebrain stimulation on cortical state (España and Berridge, 2006). Thus, combining detailed characterization of sensory and interoceptive inputs with the activity monitoring of other neuromodulatory nuclei may help isolate the specific contribution of LC bursts to electrocortical spectral changes.

Another open question is whether this LC-induced pattern of spectral changes reflects the activity of neural oscillators or aperiodic activity (also called scale-free or fractal activity) (He, 2014). A stimulus-driven decrease in the aperiodic spectral exponent (i.e., a slower decay of power spectral density with frequency) could result in similar increases of high-frequency and decreases of low-frequency activity, consistent with a shift towards greater excitation relative to inhibition (Podvalny et al., 2015; Donoghue et al., 2020). We did not test this possibility, which could be explored in future studies with time-resolved spectral parameterisation using, for example, the SPRiNT extension of the Brainstorm toolbox for MATLAB (Wilson et al., 2022; Hebron et al., 2025). This answer would improve the mechanistic understanding of the global effects of phasic LC activation, and link the current results to theories of brain “criticality” (Chaudhuri et al., 2018; Beggs, 2022). Nevertheless, our main conclusion that the spectral changes we report here indicate a sustained increase in arousal driven by LC phasic activation from *∼*0.1 to 1 s is consistent with both possibilities (i.e. that LC-induced spectral changes reflect the activity of neural oscillators or aperiodic activity), given that arousal is associated with changes of the spectral exponent (Lendner et al., 2020; Hebron et al., 2025).

### LC burst contribution to the time domain VP elicited by sudden sensory events

The present results also have clear implications for the understanding of the large amplitude, time-domain VP response elicited by SISS. We have recently demonstrated the crucial role of the diffuse and non-specific extralemniscal thalamocortical sensory system in generating the VP (Somervail et al., 2025, 2021, 2022). However, it is unknown whether and to which degree the LC contributes to this widespread cortical response. Two of our current observations - that (1) the magnitude of spontaneous LC bursting predicts the amplitude of the ECoG negativity, and (2) pharmacological suppression of NA release by systemic clonidine administration reduces the amplitude of the ECoG response (i.e. the VP) elicited by auditory SISS - indicate that the NA release consequent to LC activation at least partially contribute to the recorded VP. However, the network mechanisms of the LC contribution to the VP are unclear, and our observations are consistent with at least two possible physiological pathways.

The first is that the phasic activation of the LC causes, via diffuse LC-cortical projections (Constantinople and Bruno, 2011), an electrocortical negativity that temporally coincides with the VP, but it does not directly contribute to it. This possibility would explain well why the negative ECoG deflection in response to auditory stimulation is so much larger than that associated with phasic LC activation alone. This explanation would also be consistent with the observation that both the time domain ECoG response to auditory SISS (Figure 4) and the corresponding VP recorded in human EEG (Somervail et al., 2025) consist of a positive deflection following the negativity, while the ECoG signal associated with spontaneous phasic LC activation was a unipolar negativity (Figures 1, 4, and 5). In fact, the auditory-evoked positive deflection was even more pronounced in the clonidine condition, when the LC-evoked negativity was reduced in amplitude - possibly because of the overlap between the two potentials, with the LC-induced negativity obscuring the auditory-evoked positivity.

**Figure 5.**
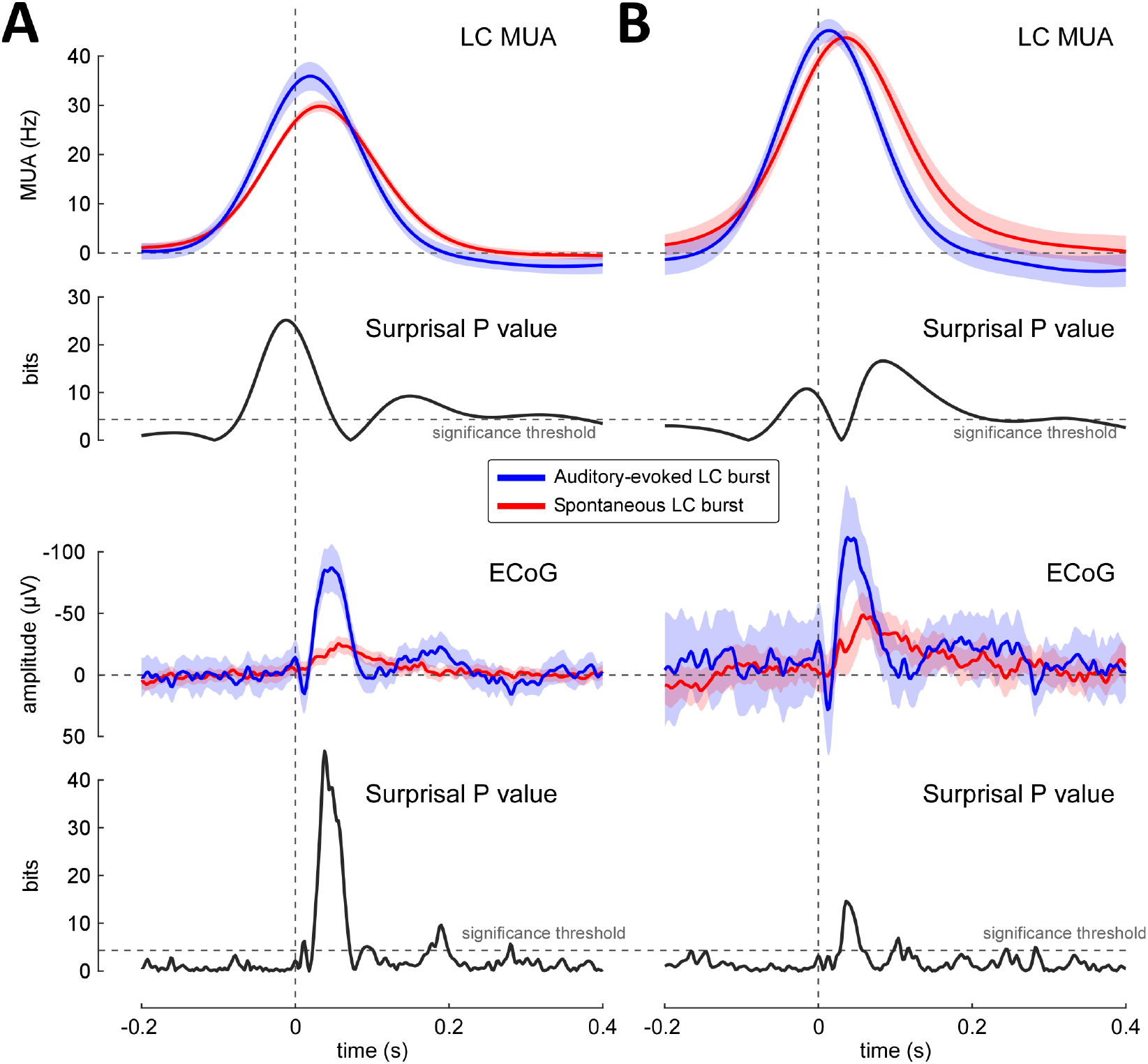
Within-subject comparison of ECoG responses associated with spontaneous and auditory-evoked LC burst. **(A)** The LC SDF (top row) and ECoG responses (third row) associated with auditory stimulation (blue) and spontaneous LC burst (red) during wakefulness (n = 1 rat). The associated surprisal P-values below each comparison show the result of a t-test between spontaneous and auditory-evoked LC bursts. Shaded areas indicate 95% confidence intervals computed with t-tests against zero. The mean LC burst frequency and ECoG response evoked by auditory stimulation were very consistently larger than the spontaneous LC bursts, with surprisal values exceeding 20 and 40 bits respectively (i.e., P - values lower than 10^-6^ and 10^-13^ respectively). **(B)** To test whether the difference in LC burst frequency accounted for the difference in ECoG response, we selected a subset of LC bursts from each condition with comparable peak magnitudes. The resulting ECoG responses to auditory stimulation were still substantially larger than those associated with spontaneous LC bursts, but the difference was reduced and became somewhat less consistent. These results indicate that the difference in amplitude of the ECoG response associated with auditory stimuli compared to spontaneous LC burst was only partly accounted for by the higher-frequency LC burst elicited by auditory stimuli, indicating the involvement of other brain systems in the generation of the surprise response to auditory stimuli.

The second explanation is that NA release from the LC in the non-specific thalamus facilitates transmission of contextual information about the surprising event via LC-NA inputs for the extralemniscal pathway (Lindvall et al., 1974; Schwarz et al., 2015), and therefore directly contributes to the VP magnitude. This explanation is the most plausible given that LC activation is highly sensitive to the interdependent factors of novelty, surprise, and salience (Foote et al., 1980; Aston-Jones and Bloom, 1981b; Rasmussen et al., 1986; Sara et al., 1994; Hervé-Minvielle and Sara, 1995). Furthermore, LC responsiveness signals high-level contextual information, such as the behavioural relevance of a stimulus (Bouret and Sara, 2004; Edeline et al., 2011; Martins and Froemke, 2015), and LC receives widespread cortical inputs capable of carrying both sensory and high-level contextual information (Luppi et al., 1995; Schwarz et al., 2015; Breton-Provencher and Sur, 2019). In this framework, the LC activation is a key component of the extralemniscal activity resulting in the widespread cortical modulation indexed by the VP. Thus, the suggested neuromodulatory mechanism could explain how the VP magnitude is sensitive to the surprise content of the eliciting sensory stimulus (Iannetti et al., 2008; Valentini et al., 2011; Torta et al., 2012; Ronga et al., 2013; Somervail et al., 2021, 2022).

Unlike the direct application of clonidine to the LC (Neves et al., 2018), its systemic administration is unlikely to completely abolish the noradrenergic effects of phasic LC activation. Therefore, the remaining time-domain cortical responses to auditory stimulation that we observed after the systemic administration of clonidine (Figure 6) may be due to residual LC activity and NA release. Therefore, we could not discern the exact degree of contribution of the LC to the VP elicited by SISS through either of the pathways described above. These questions shall be addressed in future studies.

**Figure 6.**
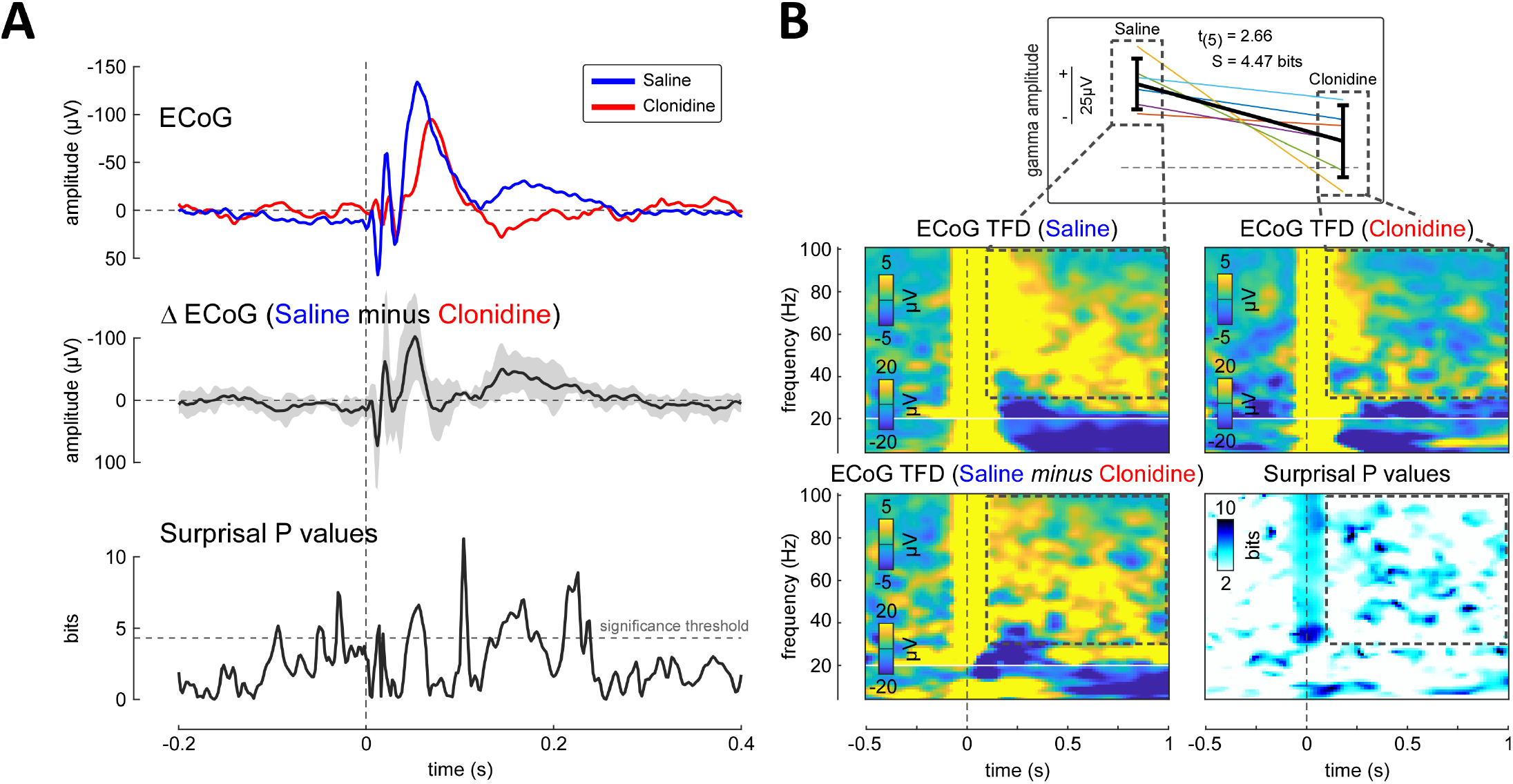
Comparison of cortical responses evoked by auditory stimulation during blockade of LC noradrenaline release by clonidine. **(A, top)** Group-level average time domain ECoG responses to auditory stimulation in the saline (blue) and clonidine (red) conditions. **(A, middle)** Average difference of the ECoG responses between the saline and clonidine conditions. Shaded area indicates 95% confidence intervals computed with point-by-point paired t-tests. **(A, bottom)** Surprisal values from the paired t-tests. The presence of clonidine reduced the amplitude of the early time-domain response and eliminated the late negative deflection at *∼*0.12 to *∼*0.22 s post-stimulus. **(B)** Group-level average time-frequency responses to auditory stimuli in the saline (*top right*) and clonidine (*top right*) conditions, together with their difference (*bottom left*) and surprisal P values (point-by-point t-test, *bottom right*). Note the two different colour scales to highlight amplitude changes of low (5 - 20 Hz) and high (20 - 100 Hz) frequencies. The top panel shows the mean amplitude values extracted in each subject from the high-frequency band (30 - 100 Hz) in a time-window from 0.1 to 1 s. Clonidine attenuated all auditory-evoked cortical responses: the transient high-frequency amplitude increase (30 - 100 Hz, around time of burst onset), the long-lasting high-frequency amplitude increase (30 - 100 Hz, from 0.1 to 1 s), and the long-lasting decrease of low-frequency amplitude (5 - 20 Hz, from *∼*0.25 to 1 s). These results suggest that noradrenaline release contributes to the transient ECoG response to surprising auditory stimuli and the associated arousal.

### Exteroceptive, interoceptive and higher-order triggers of phasic LC activation

In contrast to the slow changes of tonic LC activity in the wake-sleep cycle (Aston-Jones and Bloom, 1981a), a phasic increase of LC firing rate - a burst - is observed in response to surprising sudden events like the SISS used in the current study (Figure 4) (Aston-Jones and Bloom, 1981b; Jordan, 2024). Interestingly, phasic LC activation also occurs spontaneously during wakefulness, in the absence of experimental stimulation (Figure 1). The causes of spontaneous LC bursting are unclear, but might reflect uncontrolled inputs to the LC, such as exteroceptive inputs (such as brief tones or flashes of light) (Aston-Jones and Bloom, 1981b), interoceptive signals (such as heart beat) (Osorio-Forero et al., 2021), and top-down excitatory inputs, e.g. from the prefrontal cortex (Jodo et al., 1998) or the amygdala (Bouret et al., 2003). This view is coherent with the recently provided framework postulating that the LC operates as a “global model failure system”, with an output that scales with the global rate of prediction error or surprise signals (Jordan and Keller, 2023; Jordan, 2024).

Even in an environment that seems silent to the experimenter, large surprise-related brain responses can be elicited by ultrasonic sounds that rodents can perceive, such as those triggered by experimental machinery (Hu et al., 2015). Internal bodily signals can also be unexpected and sudden, such as a sharp and intense pain in the chest or a cardiac arrhythmia, and can therefore trigger phasic LC activation (Van Bockstaele and Aston-Jones, 1995; Osorio-Forero et al., 2021). Such exteroceptive or interoceptive signals may underlie what appears to be “spontaneous” LC phasic activation. Both can also be considered bottom-up surprise signals, as they reflect a sudden change of sensory input (the lowest level of the predictive hierarchy).

The other possible cause of “spontaneous” LC phasic activation is top-down cortical input: Top-down activity is by definition further removed from the sensory input, and is perhaps more intuitively understood at longer timescales, such as the exploration of novel environments and the associated increase of tonic LC firing (Vankov et al., 1995). However, sudden top-down signals could also elicit phasic LC activation: the sudden recall of a memory - which also causes a transient increase of pupil diameter that is likely LC-mediated (El Haj et al., 2019) - or a sudden violation of a high-level internal model of the animal and its environment (i.e. if the gradual accumulation of evidence against the model suddenly exceeds a threshold). This type of top-down effect on LC is highly plausible, given that stimulation or suppression of prefrontal cortex activity can excite or inhibit LC activity (Jodo et al., 1998).

Our findings demonstrate that the magnitude of “spontaneous” LC bursts and associated spectral changes in cortical activity are typically weaker than those evoked by the SISS we presented. This discrepancy may reflect a general bias in LC sensitivity towards external over internal (such as interoceptive and top-down) events. It is also possible that the difference simply arises because the experiments were conducted with healthy animals in a safe lab environment, resulting in minimally surprising internal signals. Since we did not record any interoceptive or top-down surprise signals, we cannot rule out that an intense internally generated surprise signal might elicit a phasic LC activation as large as that elicited by a surprising sensory stimulus.

### A brain system for rapid and reactive behavioural response to environmental surprise

Integrating these findings with our previous research into the functional significance of the non-specific VP elicited by SISS (Mouraux et al., 2003; Somervail et al., 2025) paints a picture of the overall CNS response to surprise. At cortical level, this response is characterised by at least two components: (1) a rapid (lasting 500 ms) time-domain transient primarily mediated by diffuse thalamocortical projections of the extralemniscal system and likely reflecting a ‘cortical reset’ (Perovic et al., 2025; Somervail et al., 2025), and (2) a sustained (lasting at least a few seconds) spectral shift towards higher frequencies. These two components reflect the cortical consequences of a phasic response of the LC to sudden surprise, in contrast to a slower tonic LC activation and sustained arousal associated with exploration of a novel environment (Van Bockstaele and Aston-Jones, 1995) and simulated by pharmacological modulation of LC activity (Berridge et al., 1993). This two-part phasic response may serve to suppress less urgent behaviours, reallocating neural resources to facilitate rapid execution of contextually urgent actions. This interpretation aligns well with the view that phasic LC activation results in an interruption of ongoing activity and dynamic reorganisation of brain networks to allow rapid “cognitive shifts” and behavioural adaptation (Bouret and Sara, 2005).

## Conclusion

In conclusion, these results highlight a crucial role of the LC in generating the electrophysiological response to surprising environmental events. Future research can elucidate the mechanistic relationship between the LC and other components of the extralemniscal system, such as the non-specific thalamus.

## Data availability

The data underlying this article are available on request from the corresponding authors (O.E. and G.I.).

## Author contributions statement

O.E. and M.Y. conceived the experiment, M.Y. conducted the experiment, R.S. analysed the results. R.S., G.I., and O.E. wrote and reviewed the manuscript.

## Acknowledgments

We thank Ernesto Duran for his help in chronic electrode implantation.

## Supplementary Material

**Supplementary Table 1.** The list of rats used for each experimental condition.

| Rat ID | Auditory Stimulation<sup>a</sup> | Pharmacology<sup>b</sup> | Microstimulation<sup>c</sup> | LC neuronal activity |
| --- | --- | --- | --- | --- |
| 1073 | n/a | n/a | n/a | x |
| 1405 | 40ms, 100 dB SPL | x | n/a | x |
| 1575 | n/a | n/a | n/a | x |
| 1576 | n/a | n/a | n/a | x |
| 1574 | 40ms, 90-105 dB SPL | n/a | n/a | x |
| 5794 | 60ms, 100 dB SPL | n/a | x | n/a |
| 5933 | 40ms, 90-105 dB SPL | n/a | x | n/a |
| 5934 | 60ms, 100 dB SPL | n/a | x | n/a |
| 911 | 60ms, 105 dB SPL | n/a | x | n/a |
| 912 | 60ms, 105 dB SPL | n/a | x | n/a |
| 401 | 20ms, 100 dB SPL | n/a | x | n/a |
| 402 | 20ms, 100 dB SPL | n/a | x | n/a |
| 1311 | 20ms, 100 dB SPL | x | n/a | n/a |
| 1313 | 20ms, 100 dB SPL | x | n/a | n/a |
| 1323 | 40ms, 100 dB SPL | x | n/a | n/a |
| 1401 | 40ms, 100 dB SPL | x | n/a | n/a |
| 1404 | 40ms, 100 dB SPL | x | n/a | n/a |

<sup>a</sup> - white noise (20 - 60-ms, ISI: 10 - 20 s),
<sup>b</sup> - clonidine (0.05 mg/kg, i.p.),
<sup>c</sup> - electric pulses (100 ms, 50 uA at 20, 50, 100 Hz; ISI: 10 - 15 s).

**Supplementary Table 2.** Number and percentage of rejected trials.

| Experimental Condition | Total number (MEAN ± SD) | Percent (MEAN ± SD) |
| --- | --- | --- |
| Auditory | 5 ± 17 | 7 ± 18 |
| Auditory (clonidine) | 1 ± 2 | 6 ± 14 |
| LC stimulation 20 Hz | 0 | 0 |
| LC stimulation 50 Hz | 0 | 0 |
| LC stimulation 100 Hz | 0 | 0 |
| Spontaneous LC burst | 22 ± 20 | 8 ± 10 |

## References

H. A. Abrahamian, T. Allison, W. R. Goff, and B. S. Rosner. Effects OF THIOPENTAL ON HUMAN CEREBRAL EVOKED RESPONSES. Anesthesiology, 24:650–657, 1963. ISSN 0003-3022. doi: 10.1097/00000542-196309000-00013. PMID: 14063764.

G. Aston-Jones and F. Bloom. Activity of norepinephrine-containing locus coeruleus neurons in behaving rats anticipates fluctuations in the sleep-waking cycle. The Journal of Neuroscience, 1(8):876–886, aug 1 1981a. ISSN 0270-6474, 1529-2401. doi: 10.1523/JNEUROSCI.01-08-00876.1981.

G. Aston-Jones and F. Bloom. Norepinephrine-containing locus coeruleus neurons in behaving rats exhibit pronounced responses to non-noxious environmental stimuli. The Journal of Neuroscience, 1(8):887–900, aug 1 1981b. ISSN 0270-6474, 1529-2401. doi: 10.1523/JNEUROSCI.01-08-00887.1981.

J. Bancaud, V. Bloch, and J. Paillard. [Encephalography; a study of the potentials evoked in man on the level with the vertex]. Revue neurologique, 89(5):399–418, 1953. ISSN 0035-3787. PMID: 13167923 publisher-place: France.

A. M. Bastos, J. A. Donoghue, S. L. Brincat, M. Mahnke, J. Yanar, J. Correa, A. S. Waite, M. Lundqvist, J. Roy, E. N. Brown, and E. K. Miller. Neural effects of propofol-induced unconsciousness and its reversal using thalamic stimulation. eLife, 10:1–28, 2021. ISSN 2050084X. doi: 10.7554/ELIFE.60824. PMID: 33904411.

J. M. Beggs. The cortex and the critical point: understanding the power of emergence. MIT Press, 2022. ISBN 0-262-54403-2.

M. Bellesi, B. A. Riedner, G. N. Garcia-Molina, C. Cirelli, and G. Tononi. Enhancement of sleep slow waves: Underlying mechanisms and practical consequences. Frontiers in Systems Neuroscience, 8:1–17, 2014. ISSN 16625137. doi: 10.3389/fnsys.2014.00208.

G. Berlucchi. One or many arousal systems? Reflections on some of Giuseppe Moruzzi’s foresights and insights about the intrinsic regulation of brain activity. Archives Italiennes De Biologie, 135(1):5–14, 1 1997. ISSN 0003-9829. PMID: 9139583.

C. Berridge, M. Page, R. Valentino, and S. Foote. Effects of locus coeruleus inactivation on electroencephalographic activity in neocortex and hippocampus. Neuroscience, 55(2):381–393, 7 1993. ISSN 03064522. doi: 10.1016/0306-4522(93)90507-C.

C. W. Berridge. Noradrenergic modulation of arousal. Brain Research Reviews, 58(1):1–17, 6 2008. ISSN 01650173. doi: 10.1016/j.brainresrev.2007.10.013.

S. Bouret and S. J. Sara. Reward expectation, orientation of attention and locus coeruleusmedial frontal cortex interplay during learning. European Journal of Neuroscience, 20(3):791–802, 8 2004. ISSN 0953-816X, 1460-9568. doi: 10.1111/j.1460-9568.2004.03526.x.

S. Bouret and S. J. Sara. Network reset: A simplified overarching theory of locus coeruleus noradrenaline function. Trends in Neurosciences, 28(11):574–582, 2005. ISSN 01662236. doi: 10.1016/j.tins.2005.09.002. PMID: 16165227.

S. Bouret, A. Duvel, S. Onat, and S. J. Sara. Phasic Activation of Locus Ceruleus Neurons by the Central Nucleus of the Amygdala. The Journal of Neuroscience, 23(8):3491–3497, apr 15 2003. ISSN 0270-6474, 1529-2401. doi: 10.1523/JNEUROSCI.23-08-03491.2003.

V. Breton-Provencher and M. Sur. Active control of arousal by a locus coeruleus GABAergic circuit. Nature Neuroscience, 22(2): 218–228, 2 2019. ISSN 1097-6256, 1546-1726. doi: 10.1038/s41593-018-0305-z.

R. Chaudhuri, B. J. He, and X.-J. Wang. Random Recurrent Networks Near Criticality Capture the Broadband Power Distribution of Human ECoG Dynamics. Cerebral Cortex, 28(10):3610–3622, oct 1 2018. ISSN 1047-3211, 1460-2199. doi: 10.1093/cercor/bhx233.

C. M. Constantinople and R. M. Bruno. Effects and Mechanisms of Wakefulness on Local Cortical Networks. Neuron, 69(6):1061–1068, 3 2011. ISSN 08966273. doi: 10.1016/j.neuron.2011.02.040.

H. Davis and S. Zerlin. Acoustic Relations of the Human Vertex Potential. The Journal of the Acoustical Society of America, 39(1): 109–116, 1 1966. ISSN 0001-4966. doi: 10.1121/1.1909858.

P. A. Davis. Effects OF ACOUSTIC STIMULI ON THE WAKING HUMAN BRAIN. Journal of Neurophysiology, 2(6):494–499, nov 1 1939. ISSN 0022-3077. doi: 10.1152/jn.1939.2.6.494.

J. W. De Gee, K. Tsetsos, L. Schwabe, A. E. Urai, D. McCormick, M. J. McGinley, and T. H. Donner. Pupil-linked phasic arousal predicts a reduction of choice bias across species and decision domains. eLife, 9:e54014, jun 16 2020. ISSN 2050-084X. doi: 10.7554/eLife.54014.

A. Delorme and S. Makeig. Eeglab: an open source toolbox for analysis of single-trial EEG dynamics. Journal of Neuroscience Methods, 13:9–21, 2004. ISSN 0165-0270. doi: 10.1016/j.jneumeth.2003.10.009. PMID: 15102499 arXiv: 10.1016/j.jneumeth.2003.10.009. ISBN: 0165-0270 (Print)\r0165-0270 (Linking).

T. Donoghue, M. Haller, E. J. Peterson, P. Varma, P. Sebastian, R. Gao, T. Noto, A. H. Lara, J. D. Wallis, R. T. Knight, A. Shestyuk, and B. Voytek. Parameterizing neural power spectra into periodic and aperiodic components. Nature Neuroscience, 23(12):1655–1665, 12 2020. ISSN 1097-6256, 1546-1726. doi: 10.1038/s41593-020-00744-x.

H. Dringenberg and M. Olmstead. Integrated contributions of basal forebrain and thalamus to neocortical activation elicited by pedunculopontine tegmental stimulation in urethane-anesthetized rats. Neuroscience, 119(3):839–853, 7 2003. ISSN 03064522. doi: 10.1016/S0306-4522(03)00197-0.

J.-M. Edeline, Y. Manunta, and E. Hennevin. Induction of selective plasticity in the frequency tuning of auditory cortex and auditory thalamus neurons by locus coeruleus stimulation. Hearing Research, 274(1-2):75–84, 4 2011. ISSN 03785955. doi: 10.1016/j.heares.2010.08.005.

M. El Haj, S. M. J. Janssen, K. Gallouj, and Q. Lenoble. Autobiographical memory increases pupil dilation. Translational Neuroscience, 10(1):280–287, dec 19 2019. ISSN 2081-6936. doi: 10.1515/tnsci-2019-0044.

O. Eschenko, C. Magri, S. Panzeri, and S. J. Sara. Noradrenergic neurons of the locus coeruleus are phase locked to cortical up-down states during sleep. Cerebral Cortex, 22(2):426–435, 2012. ISSN 10473211. doi: 10.1093/cercor/bhr121. PMID: 21670101.

R. A. España and C. W. Berridge. Organization of noradrenergic efferents to arousalrelated basal forebrain structures. Journal of Comparative Neurology, 496(5):668–683, jun 10 2006. ISSN 0021-9967, 1096-9861. doi: 10.1002/cne.20946.

S. L. Foote, G. Aston-Jones, and F. E. Bloom. Impulse activity of locus coeruleus neurons in awake rats and monkeys is a function of sensory stimulation and arousal. Proceedings of the National Academy of Sciences, 77(5):3033–3037, 5 1980. ISSN 0027-8424, 1091-6490. doi: 10.1073/pnas.77.5.3033.

E. Glennon, I. Carcea, A. R. O. Martins, J. Multani, I. Shehu, M. A. Svirsky, and R. C. Froemke. Locus coeruleus activation accelerates perceptual learning. Brain Research, 1709:39–49, 4 2019. ISSN 00068993. doi: 10.1016/j.brainres.2018.05.048.

M. Goard and Y. Dan. Basal forebrain activation enhances cortical coding of natural scenes. Nature Neuroscience, 12(11):1444–1449, 11 2009. ISSN 1097-6256, 1546-1726. doi: 10.1038/nn.2402.

S. Greenland. Invited Commentary: The Need for Cognitive Science in Methodology. American Journal of Epidemiology, 186(6): 639–645, sep 15 2017. ISSN 0002-9262, 1476-6256. doi: 10.1093/aje/kwx259.

B. J. He. Scale-free brain activity: past, present, and future. Trends in Cognitive Sciences, 18(9):480–487, 9 2014. ISSN 13646613. doi: 10.1016/j.tics.2014.04.003.

H. Hebron, A. M. Stephan, J. Cataldi, and F. Siclari. Aperiodic eeg activity provides a linear, bidirectional, and spatially uniform marker of subjective and objective vigilance in humans, both within and across states. bioRxiv, 2025. doi: 10.1101/2025.08.30.673229. URL https://www.biorxiv.org/content/early/2025/09/02/2025.08.30.673229.

A. Hervé-Minvielle and S. J. Sara. Rapid habituation of auditory responses of locus coeruleus cells in anaesthetized and awake rats:. NeuroReport, 6(10):1363–1368, 7 1995. ISSN 0959-4965. doi: 10.1097/00001756-199507100-00001.

L. Hu and G. D. Iannetti. Neural indicators of perceptual variability of pain across species. Proceedings of the National Academy of Sciences, 116(5):1782–1791, jan 29 2019. ISSN 1091-6490. doi: 10.1073/pnas.1812499116. PMID: 30642968 ISBN: 1812499116.

L. Hu, X. L. Xia, W. W. Peng, W. X. Su, F. Luo, H. Yuan, A. T. Chen, M. Liang, and G. Iannetti. Was it a pain or a sound? Across-species variability in sensory sensitivity. Pain, 156(12):2449–57, 2015. ISSN 1872-6623. doi: 10.1097/j.pain.0000000000000316. PMID: 26270592 ISBN: 0000000000000.

J. Hunter and H. H. Jasper. Effects of thalamic stimulation in unanaesthetised animals. The arrest reaction and petit Mal-like seizures, activation patterns and generalized convulsions. Electroencephalography and Clinical Neurophysiology, 1(1-4):305–324, 1949. ISSN 00134694. doi: 10.1016/0013-4694(49)90196-0. PMID: 18135423.

L. Hurley, D. Devilbiss, and B. Waterhouse. A matter of focus: monoaminergic modulation of stimulus coding in mammalian sensory networks. Current Opinion in Neurobiology, 14(4):488–495, 8 2004. ISSN 09594388. doi: 10.1016/j.conb.2004.06.007.

G. D. Iannetti, N. P. Hughes, M. C. Lee, and A. Mouraux. Determinants of laser-evoked EEG responses: pain perception or stimulus saliency? Journal of Neurophysiology, 100(2):815–828, 2008. ISSN 0022-3077. doi: 10.1152/jn.00097.2008. PMID: 18525021 ISBN: 0022-3077 (Print) \r0022-3077 (Linking).

H. H. Jasper. Diffuse projection systems: The integrative action of the thalamic reticular system. Electroencephalography and Clinical Neurophysiology, 1(1-4):405–420, 1949. ISSN 00134694. doi: 10.1016/0013-4694(49)90213-8. PMID: 18421831.

E. Jodo, C. Chiang, and G. Aston-Jones. Potent excitatory influence of prefrontal cortex activity on noradrenergic locus coeruleus neurons. Neuroscience, 83(1):63–79, 1998. ISSN 0306-4522. doi: 10.1016/s0306-4522(97)00372-2. publisher: Elsevier BV.

B. E. Jones. Arousal systems. Frontiers in Bioscience, 8(6):s438–451, 2003. ISSN 10939946. doi: 10.2741/1074. PMID: 12700104.

R. Jordan. The locus coeruleus as a global model failure system. Trends in Neurosciences, 47(2):92–105, 2024. ISSN 1878108X. doi: 10.1016/j.tins.2023.11.006. PMID: 38102059 publisher: The Author.

R. Jordan and G. B. Keller. The locus coeruleus broadcasts prediction errors across the cortex to promote sensorimotor plasticity. eLife, 12:RP85111, jun 7 2023. ISSN 2050-084X. doi: 10.7554/eLife.85111.

J. D. Lendner, R. F. Helfrich, B. A. Mander, L. Romundstad, J. J. Lin, M. P. Walker, P. G. Larsson, and R. T. Knight. An electrophysiological marker of arousal level in humans. eLife, 9:e55092, jul 28 2020. ISSN 2050-084X. doi: 10.7554/eLife.55092.

S.-C. Lin, D. Gervasoni, and M. A. L. Nicolelis. Fast Modulation of Prefrontal Cortex Activity by Basal Forebrain Noncholinergic Neuronal Ensembles. Journal of Neurophysiology, 96(6):3209–3219, 12 2006. ISSN 0022-3077, 1522-1598. doi: 10.1152/jn.00524.2006.

O. Lindvall, A. Björklund, A. Nobin, and U. Stenevi. The adrenergic innervation of the rat thalamus as revealed by the glyoxylic acid fluorescence method. Journal of Comparative Neurology, 154(3):317–347, 4 1974. ISSN 0021-9967, 1096-9861. doi: 10.1002/cne.901540307.

P.-H. Luppi, G. Aston-Jones, H. Akaoka, G. Chouvet, and M. Jouvet. Afferent projections to the rat locus coeruleus demonstrated by retrograde and anterograde tracing with cholera-toxin B subunit and Phaseolus vulgaris leucoagglutinin. Neuroscience, 65(1): 119–160, 3 1995. ISSN 03064522. doi: 10.1016/0306-4522(94)00481-J.

A. R. O. Martins and R. C. Froemke. Coordinated forms of noradrenergic plasticity in the locus coeruleus and primary auditory cortex. Nature Neuroscience, 18(10):1483–1492, 10 2015. ISSN 1097-6256, 1546-1726. doi: 10.1038/nn.4090.

A. Marzo, N. K. Totah, R. M. Neves, N. K. Logothetis, and O. Eschenko. Unilateral electrical stimulation of rat locus coeruleus elicits bilateral response of norepinephrine neurons and sustained activation of medial prefrontal cortex. Journal of Neurophysiology, 111 (12):2570–2588, jun 15 2014. ISSN 0022-3077, 1522-1598. doi: 10.1152/jn.00920.2013.

J. McBurney-Lin, Y. Sun, L. S. Tortorelli, Q. A. T. Nguyen, S. Haga-Yamanaka, and H. Yang. Bidirectional pharmacological perturbations of the noradrenergic system differentially affect tactile detection. Neuropharmacology, 174:108151, 9 2020. ISSN 00283908. doi: 10.1016/j.neuropharm.2020.108151.

M. Moayedi, G. Di Stefano, M. T. Stubbs, B. Djeugam, M. Liang, and G. D. Iannetti. Nociceptive-evoked potentials are sensitive to behaviorally relevant stimulus displacements in egocentric coordinates. eNeuro, 3(3):399–418, 2016. ISSN 23732822. doi: 10.1523/ENEURO.0151-15.2016. PMID: 27419217.

A. Mouraux and G. D. Iannetti. Nociceptive Laser-Evoked Brain Potentials Do Not Reflect Nociceptive-Specific Neural Activity. Journal of neurophysiology, 101(6):3258–69, 6 2009. ISSN 0022-3077. doi: 10.1152/jn.91181.2008. PMID: 19339457 ISBN: 0022-3077 (Print) 0022-3077 (Linking).

A. Mouraux, J. Guérit, and L. Plaghki. Non-phase locked electroencephalogram (EEG) responses to CO2 laser skin stimulations may reflect central interactions between A- and C-fibre afferent volleys. Clinical Neurophysiology, 114(4):710–722, 4 2003. ISSN 13882457. doi: 10.1016/S1388-2457(03)00027-0.

R. Munn, E. J. Müller, G. Wainstein, and J. M. Shine. The ascending arousal system shapes neural dynamics to mediate awareness of cognitive states. Nature Communications, 12(1):6016, oct 14 2021. ISSN 2041-1723. doi: 10.1038/s41467-021-26268-x.

R. M. Neves, S. Van Keulen, M. Yang, N. K. Logothetis, and O. Eschenko. Locus coeruleus phasic discharge is essential for stimulus-induced gamma oscillations in the prefrontal cortex. Journal of Neurophysiology, 119(3):904–920, mar 1 2018. ISSN 0022-3077, 1522-1598. doi: 10.1152/jn.00552.2017.

G. Novembre, V. M. Pawar, R. J. Bufacchi, M. Kilintari, M. Srinivasan, J. C. Rothwell, P. Haggard, and G. D. Iannetti. Saliency detection as a reactive process: Unexpected Sensory Events Evoke Corticomuscular Coupling. The Journal of Neuroscience, 38(9): 2385–2397, 2018. ISSN 0270-6474. doi: 10.1523/JNEUROSCI.2474-17.2017. PMID: 29378865.

Y. Novitskaya, S. J. Sara, N. K. Logothetis, and O. Eschenko. Ripple-triggered stimulation of the locus coeruleus during post-learning sleep disrupts ripple/spindle coupling and impairs memory consolidation. Learning and Memory, 23(5):238–248, 2016. ISSN 15495485. doi: 10.1101/lm.040923.115. PMID: 27084931.

A. Nuñez. Unit activity of rat basal forebrain neurons: Relationship to cortical activity. Neuroscience, 72(3):757–766, 6 1996. ISSN 03064522. doi: 10.1016/0306-4522(95)00582-X.

A. Osorio-Forero, R. Cardis, G. Vantomme, A. Guillaume-Gentil, G. Katsioudi, C. Devenoges, L. M. Fernandez, and A. Lüthi. Noradrenergic circuit control of non-REM sleep substates. Current Biology, 31(22):5009–5023.e7, 11 2021. ISSN 09609822. doi: 10.1016/j.cub.2021.09.041.

W. Peng, X. Xia, M. Yi, G. Huang, Z. Zhang, G. Iannetti, and L. Hu. Brain oscillations reflecting pain-related behavior in freely moving rats. Pain, 159(1):106–118, 2018. ISSN 0304-3959 (Print) 0304-3959. doi: 10.1097/j.pain.0000000000001069.1872-6623 Peng, Weiwei Xia, Xiaolei Yi, Ming Huang, Gan Zhang, Zhiguo Iannetti, Giandomenico Hu, Li PAIN JLARAXR/Wellcome Trust/United Kingdom Journal Article Research Support, Non-U.S. Gov’t United States 2017/09/28 Pain. 2018 Jan;159(1):106–118. doi: 10.1097/j.pain.0000000000001069.

S. Perovic, R. Somervail, D. Benusiglio, and G. D. Iannetti. The Extralemniscal System Modulates Early Somatosensory Cortical Processing. The Journal of Neuroscience, 45(40), 2025. ISSN 0270-6474, 1529-2401. doi: 10.1523/JNEUROSCI.1815-24.2025. URL https://www.jneurosci.org/lookup/doi/10.1523/JNEUROSCI.1815-24.2025. [Online; accessed 2025-10-15].

E. Podvalny, N. Noy, M. Harel, S. Bickel, G. Chechik, C. E. Schroeder, A. D. Mehta, M. Tsodyks, and R. Malach. A unifying principle underlying the extracellular field potential spectral responses in the human cortex. Journal of Neurophysiology, 114(1):505–519, 7 2015. ISSN 0022-3077, 1522-1598. doi: 10.1152/jn.00943.2014.

K. Rasmussen, D. A. Morilak, and B. L. Jacobs. Single unit activity of locus coeruleus neurons in the freely moving cat. Brain Research, 371(2):324–334, 4 1986. ISSN 00068993. doi: 10.1016/0006-8993(86)90370-7.

M. J. Redinbaugh, J. M. Phillips, N. A. Kambi, S. Mohanta, S. Andryk, G. L. Dooley, M. Afrasiabi, A. Raz, and Y. B. Saalmann. Thalamus Modulates Consciousness via Layer-Specific Control of Cortex. Neuron, 106(1):66–75.e12, 2020. ISSN 10974199. doi: 10.1016/j.neuron.2020.01.005. PMID: 32053769 publisher: Elsevier Inc.

I. Ronga, E. Valentini, A. Mouraux, and G. D. Iannetti. Novelty is not enough: laser-evoked potentials are determined by stimulus saliency, not absolute novelty. Journal of neurophysiology, 44(3):692–701, 2013. ISSN 1522-1598. doi: 10.1152/jn.00464.2012. PMID: 23136349 ISBN: 0022-3077.

J. A. Ross and E. J. Van Bockstaele. The Locus Coeruleus-Norepinephrine System in Stress and Arousal: Unraveling Historical, Current, and Future Perspectives. Frontiers in Psychiatry, 11:601519, jan 27 2021. ISSN 1664-0640. doi: 10.3389/fpsyt.2020.601519.

S. J. Sara and S. Bouret. Orienting and Reorienting: The Locus Coeruleus Mediates Cognition through Arousal. Neuron, 76(1):130–141, 10 2012. ISSN 08966273. doi: 10.1016/j.neuron.2012.09.011.

S. J. Sara, A. Vankov, and A. Hervé. Locus coeruleus-evoked responses in behaving rats: A clue to the role of noradrenaline in memory. Brain Research Bulletin, 35(5-6):457–465, 1 1994. ISSN 03619230. doi: 10.1016/0361-9230(94)90159-7.

L. A. Schwarz, K. Miyamichi, X. J. Gao, K. T. Beier, B. Weissbourd, K. E. DeLoach, J. Ren, S. Ibanes, R. C. Malenka, E. J. Kremer, and L. Luo. Viral-genetic tracing of the input–output organization of a central noradrenaline circuit. Nature, 524(7563):88–92, aug 6 2015. ISSN 0028-0836, 1476-4687. doi: 10.1038/nature14600.

R. Somervail, F. Zhang, G. Novembre, R. J. Bufacchi, Y. Guo, M. Crepaldi, L. Hu, and G. D. Iannetti. Waves of Change: Brain Sensitivity to Differential, not Absolute, Stimulus Intensity is Conserved Across Humans and Rats. Cerebral Cortex, 31(2):949–960, jan 5 2021. ISSN 1047-3211. doi: 10.1093/cercor/bhaa267.

R. Somervail, R. J. Bufacchi, C. Salvatori, L. Neary-Zajiczek, Y. Guo, G. Novembre, and G. D. Iannetti. Brain Responses to Surprising Stimulus Offsets: Phenomenology and Functional Significance. Cerebral Cortex, 32(10):2231–2244, may 14 2022. ISSN 1047-3211. doi: 10.1093/cercor/bhab352.

R. Somervail, S. Perovic, R. J. Bufacchi, R. Caminiti, and G. D. Iannetti. A two-system theory of sensory-evoked brain responses. Brain, 2025. ISSN 0006-8950. doi: 10.1093/brain/awaf402. URL https://doi.org/10.1093/brain/awaf402.

J. Tasserie, L. Uhrig, J. D. Sitt, D. Manasova, M. Dupont, S. Dehaene, and B. Jarraya. Deep brain stimulation of the thalamus restores signatures of consciousness in a nonhuman primate model. Science Advances, 8(11):1–18, 2022. ISSN 23752548. doi: 10.1126/sciadv.abl5547. PMID: 35302854.

L. Tiemann, E. Schulz, J. Gross, and M. Ploner. Gamma oscillations as a neuronal correlate of the attentional effects of pain. Pain, 150 (2):302–308, 8 2010. ISSN 0304-3959. doi: 10.1016/j.pain.2010.05.014.

D. M. Torta, M. Liang, E. Valentini, A. Mouraux, and G. D. Iannetti. Dishabituation of laser-evoked EEG responses: Dissecting the effect of certain and uncertain changes in stimulus spatial location. Experimental Brain Research, 218(3):361–372, 2012. ISSN 00144819. doi: 10.1007/s00221-012-3019-6. PMID: 21265604 ISBN: 1530-8898 (Electronic)\r0898-929X (Linking).

N. K. Totah, N. K. Logothetis, and O. Eschenko. Synchronous spiking associated with prefrontal high γ oscillations evokes a 5-Hz rhythmic modulation of spiking in locus coeruleus. Journal of Neurophysiology, 125(4):1191–1201, apr 1 2021. ISSN 0022-3077, 1522-1598. doi: 10.1152/jn.00677.2020.

E. Valentini, D. M. E. Torta, A. Mouraux, and G. D. Iannetti. Dishabituation of Laser-evoked EEG Responses: Dissecting the Effect of Certain and Uncertain Changes in Stimulus Modality. Journal of Cognitive Neuroscience, 23(10):2822–2837, 2011. ISSN 0898-929X. doi: 10.1162/jocn.2011.21609.

E. J. Van Bockstaele and G. Aston-Jones. Integration in the Ventral Medulla and Coordination of Sympathetic, Pain and Arousal Functions. Clinical and Experimental Hypertension, 17(1-2):153–165, 1 1995. ISSN 1064-1963, 1525-6006. doi: 10.3109/10641969509087062.

A. Vankov, A. HervéMinvielle, and S. J. Sara. Response to Novelty and its Rapid Habituation in Locus Coeruleus Neurons of the Freely Exploring Rat. European Journal of Neuroscience, 7(6):1180–1187, 6 1995. ISSN 0953-816X, 1460-9568. doi: 10.1111/j.1460-9568.1995.tb01108.x.

X. Wang, Z. Li, X. Wang, J. Chen, Z. Guo, B. Qiao, and L. Qin. Effects of Phasic Activation of Locus Ceruleus on Cortical Neural Activity and Auditory Discrimination Behavior. The Journal of Neuroscience, 44(37):e1296232024, sep 11 2024. ISSN 0270-6474, 1529-2401. doi: 10.1523/JNEUROSCI.1296-23.2024.

C. J. Whyte, M. J. Redinbaugh, J. M. Shine, and Y. B. Saalmann. Thalamic contributions to the state and contents of consciousness. Neuron, 112(10):1611–1625, 2024. ISSN 10974199. doi: 10.1016/j.neuron.2024.04.019. publisher: The Authors.

L. E. Wilson, J. Da Silva Castanheira, and S. Baillet. Time-resolved parameterization of aperiodic and periodic brain activity. eLife, 11:e77348, sep 12 2022. ISSN 2050-084X. doi: 10.7554/eLife.77348.

M. Yang and O. Eschenko. Differential locus coeruleus–hippocampus interactions during offline states. bioRxiv, page 2025.09.18.677005, 2025. doi: 10.1101/2025.09.18.677005. URL https://www.biorxiv.org/content/biorxiv/early/2025/09/19/2025.09.18.677005.full.pdf.

M. Yang, N. K. Logothetis, and O. Eschenko. Phasic activation of the locus coeruleus attenuates the acoustic startle response by increasing cortical arousal. Scientific Reports, 11(1), 2021. ISSN 20452322. doi: 10.1038/s41598-020-80703-5. PMID: 33446792.

Z. Zhang, Y. Huang, X. Chen, J. Li, Y. Yang, L. Lv, J. Wang, M. Wang, Y. Wang, and Z. Wang. Statespecific Regulation of Electrical Stimulation in the Intralaminar Thalamus of Macaque Monkeys: Network and Transcriptional Insights into Arousal. Advanced Science, 11(33):2402718, 9 2024. ISSN 2198-3844, 2198-3844. doi: 10.1002/advs.202402718.

Z. G. Zhang, L. Hu, Y. S. Hung, A. Mouraux, and G. D. Iannetti. Gamma-Band Oscillations in the Primary Somatosensory Cortex - A Direct and Obligatory Correlate of Subjective Pain Intensity. Journal of Neuroscience, 32(22):7429–7438, 2012. ISSN 0270-6474. doi: 10.1523/JNEUROSCI.5877-11.2012. PMID: 22649223 ISBN: 1529-2401 (Electronic)\r0270-6474 (Linking).

